# MetaProClust-MS1: A tool for clustering metaproteomes using rapid MS1 profiling

**DOI:** 10.1101/2021.03.31.437863

**Authors:** Caitlin M.A. Simopoulos, Zhibin Ning, Leyuan Li, Mona M. Khamis, Xu Zhang, Mathieu Lavallée-Adam, Daniel Figeys

**Affiliations:** Ottawa Institute of Systems Biology and Department of Biochemistry, Microbiology and Immunology, Faculty of Medicine, University of Ottawa, Ottawa, ON, Canada; SIMM-University of Ottawa Joint Research Center in Systems and Personalized Pharmacology, University of Ottawa, Ottawa, ON, Canada

**Keywords:** Microbiome, Mass spectrometry, Metaproteomics, MS1-only, Clustering, Machine learning, Drug screening, Proteomics, Bioinformatics, Unsupervised learning

## Abstract

Metaproteomics is used to explore the composition, dynamics and function of microbial communities. However, acquiring data by tandem mass spectrometry is time consuming and resource intensive. To mediate this challenge, we present MetaProClust-MS1, a computational framework for microbiome screening developed to reduce the time required for data acquisition by mass spectrometry. In this proof-of-concept study, we tested MetaProClust-MS1 on data acquired using short 15 minute MS1-only mass spectrometry gradients and compared the results to those produced using data acquired by a traditional tandem mass spectrometry approach. MetaProClust-MS1 identified robust microbiome shifts caused by xenobiotics in both datasets. Cluster topologies were also significantly correlated. We demonstrate that MetaProClust-MS1 is able to rapidly screen microbiomes using only short MS1 profiles. This approach can be used to prioritize samples for deep metaproteomic analysis and will be especially useful in large-scale metaproteomic screens or in clinical settings where rapid results are required.

## 1 Background

Studying community dynamics using metaproteoemics has recently become more common due to its ability to evaluate both the taxonomic and functional composition of samples (Zhang et al., 2019). In a recent perspective, Kleiner (2019) describes the strengths of studying the metaproteome and the abundance of questions that can only be answered through metaproteomic technology. For example, through metaproteomics we can understand molecular and cellular phenotypes of microbial communities, the energy sources of individuals in a community (Bryson et al., 2016), and the functional changes introduced through post-translational modifications (Cheng et al., 2020).

One area where metaproteomics is particularly useful is in the study of human gut microbiomes and their connections to human health and disease. There is growing evidence to suggest that gut microbiome compositional and functional dysbiosis is driving human disease as observed in inflammatory bowel disease (IBD) (Morgan et al., 2012; Zhang et al., 2018), asthma (Arrieta et al., 2015), multiple sclerosis (Jangi et al., 2016), obesity (Sonnenburg et al., 2016), type II diabetes and cardiovascular diseases (Tang et al., 2017). In addition, xenobiotics can cause significant impacts to gut microbiomes (Maier et al., 2018) warranting further investigation into how compounds may affect the gut microbiota. Recently, Li et al. (2020c) introduced RapidAIM, an *in vitro* assay used to assess the microbiome’s response to drugs using high-throughput metaproteomics and metagenomics. Thus far, RapidAIM has been used to identify drugs that cause significant alterations to the gut microbiome (Li et al., 2020c), determine how structural analogs affect the gut microbiome (Li et al., 2020b), and explore the ways individual microbiome profiles vary in response to resistant starches (Li et al., 2020a).

Using metaproteomics as a first pass screen of samples for high-throughput screening is slow and expensive. For example, deep metaproteomics, which is used for microbial strain-level data resolution, requires upwards of 260 minute tandem mass spectrometry (MS) gradients (Zhang et al., 2017). In addition, microbiome data can be challenging to analyze due to the large amount of compositional variability observed between individuals (Schloissnig et al., 2013). Stemming from taxonomical variability, the effects of xenobiotics on gut microbiomes also have large variation when tested on multiple microbiomes (Li et al., 2020c), thus it’s essential to identify robust effects when considering high-throughput studies.

Large interest in high-throughput microbiome experiments necessitate research into reducing data acquisition time requirements. One such strategy is MS2-independent proteomics which is founded on peptide mass fingerprinting (James et al., 1993; Mann et al., 1993; Pappin et al., 1993; Yeats et al., 1993). Workflows for short gradient MS1-only proteomics, like DirectMS1 (Ivanov et al., 2020), offer the ability to identify proteins from retention time and mass-to-charge ratios of MS1 features. However, DirectMS1 remains suitable for single-species samples and has only been thoroughly tested on simple cell line samples. Unfortunately, microbiome metaproteomic samples are more complex due to the presence of multiple microbial species and contain eukaryotic host proteins that may be of interest. In addition, the common approach to metaproteomics, data-dependent mass spectrometry, acquires MS2 spectra for peptide identification based on precursor ion intensity. MS1-only data, or MS2-independent data, is not limited to these high intensity features that are likely to be peptides and thus may also contain metabolites. As such, novel computational approaches are required for data analysis of MS1-only metaproteomic data, particularly for large-scale high-throughput studies.

In this manuscript we introduce MetaProClust-MS1 (MPC-MS1), a bioinformatic tool developed for screening of high throughput metaproteomic studies through clustering experimental conditions. As proof of concept we demonstrate the ability of MPC-MS1 with a dataset of gut microbiome treated with five drugs at three different concentrations. Using a robust independent component analysis (ICA) implementation, MPC-MS1 uses k-medoids clustering to identify modules of MS1 features with similar intensity patterns. An eigenfeature vector is then calculated for each feature module and describes the intensity patterns of all features in each module. Microbiome treatments are correlated with each eigenfeature and are clustered for a quick screen of treatment effects on the microbiome. We see similar clusters identified by MPC-MS1 using MS1-only data to those observed with MS/MS acquired data indicating that our MS1-only approach with MPC-MS1 offers a rapid alternative to typical bottom-up metaprotoemics. In combination with the RapidAIM assay (Li et al., 2020c), MPC-MS1 greatly accelerates high-throughput metaproteomic studies and may be particularly useful in clinical settings.

## 2 Methods

### 2.1 Sample preparation and data acquisition

An *in vitro* RapidAIM assay (Li et al., 2020c) was used to assess the contributions of five drugs, azathioprine (AZ), ciprofloxacin (CP), diclofenac (DC), nizatidine (NZ) and paracetamol (PR) at three concentrations:high (H), medium (M) and low (L) on a human gut microbiome (fecal) sample (Table 1). The high (H) concentrations were previously identified by Li et al. (2020c) to have robust effects on a gut microbiome. Medium (M) and low (L) concentrations were used to test sensitivity of the MPC-MS1 pipeline. Using RapidAIM, a gut microbiome (fecal sample) from an individual was cultured for 48 hours in anaerobic conditions with either control treatment, dimethyl sulfoxide (DMSO), or drug treatment. Control and drug treatments at each concentration were cultured in replicates of three. Proteins were extracted from cultured samples and digested with trypsin (Worthington Biochemical Corp., Lakewood, NJ). The human stool sampling protocol (Protocol 20160585-01H) was approved by the Ottawa Health Science Network Research Ethics Board at the Ottawa Hospital.

**Table 1:** Drug concentrations used in RapidAIM treatment of gut microbiome (fecal) samples. Each drug was used at three different concentrations: low (L), medium (M) and high (H). High concentrations were those previously found to have an effect on the gut microbiome using RapidAIM as found by Li et al. (2020c).

| Drug | Concentration (μM) | | |
| --- | --- | --- | --- |
|  | Low (L) | Medium (M) | High (H) |
| Azathioprine (AZ) | 100 | 500 | 2700 |
| Ciprofloxacin (CP) | 100 | 500 | 1100 |
| Diclofenac (DC) | 100 | 500 | 3100 |
| Nizatidine (NZ) | 100 | 500 | 4500 |
| Paracetamol (PR) | 100 | 500 | 13200 |

#### 2.1.1 MS1-only data acquisition

The LC-MS analysis was performed on a Q Exactive with Easy-nLC 1200. All 48 samples samples were run in a randomized order. Peptides were separated on a tip column (75 μm inner diameter × 10 cm) packed with reverse phase beads (3 μm/120 Å ReproSil-Pur C18 resin, Dr. Maisch GmbH, Ammerbuch, Germany). 0.1% (v/v) formic acid in water was used as buffer A, and 0.1% FA in 80% acetonitrile was used as buffer B. A 15 min LC gradient was used with 4% 35% in 9min, 35% 80% in 2min, and 80% in 4min. The MS1-only method has full scans with a resolution of 70000, scan range from 200 to 1400 m/z, AGC target as 3E6, and Maximum IT as 200ms, no dependent scans followed.

#### 2.1.2 MS/MS data acquisition

The same set of digests were analyzed by LC-MS/MS and performed using an UltiMate 3000 RSLCnano system coupled to an Orbitrap Exploris 480 mass spectrometer using similar chromatographic conditions on a on a tip column (75 μm inner diameter × 20 cm) packed with reverse phase beads (3 μm/120 Å ReproSil-Pur C18 resin, Dr. Maisch GmbH, Ammerbuch, Germany). A 60-min gradient of 5 to 35% (v/v) buffer B at a 300 L/min flow rate was used. The MS full scan was performed from 350 - 1400 m/z, recorded in profile mode, and was followed by data-dependent MS/MS scan of the 12 most intense ions, a dynamic exclusion repeat count of one, and exclusion duration of 30 s. The resolutions for MS and MS/MS were 60,000 and 15,000, respectively.

### 2.2 Feature identification

#### 2.2.1 MS1-only dataset

MS1 level peptide features were identified using the OpenMS suite of tools for MS data analysis (Röst et al., 2016) with exact commands and parameters used available on the MPC-MS1 GitHub repository at https://github.com/northomics/MetaProClust-MS1/blob/main/bin/openMS_ms1_commands.sh. A brief overview of the OpenMS workflow is as follows. Thermo RAW files were first converted to ·mzML using MSConvertGUI from ProteoWizard Toolkit (Chambers et al., 2012). Peak lists were identified using PeakPickerHiRes (using default parameters other than `algorithm:signal_to_noise 0 and algorithm:ms_levels 1)`. Features output as ·XML files were identified from peaks using `FeatureFinderCentroided`. Retention times were aligned between MS runs using `MapAlignerPoseClustering` considering a maximum retention time difference of 300.0 secs and maximum m/z difference of 20.0 ppm. Finally, an experiment specific consensus map was identified using `FeatureLinkerUnlabeledQT` and the same maximum retention time and m/z differences of previously men-tioned. Linked features were then written to a ·CSV using `TextExporter` for further analysis using the MPC-MS1 workflow.

#### 2.2.2 MS/MS dataset

Spectra search and peptide quantitation were completed using MetaLab 2.0 (Cheng et al., 2020) and a closed database search of the Integrated human gut microbial Gene Catalog (IGC) (Li et al., 2014) with a peptide FDR threshold of 0.01. Peptide-level intensities were used for further analyses to maintain similarity with MS1-only features.

### 2.3 Data preprocessing

#### 2.3.1 MS1-only data

MS1 feature intensities from the OpenMS workflow and experiment metadata were imported into R v4.0.4 (R Development Core Team, 2021). First, MS1 feature intensities were divided into quartiles (Figure S1). Features belonging to the lowest quartile (intensity below 5,858,867) were considered as missing values for data filtering and only features quantified in at least 50% of each drug treatment, including the DMSO control, were kept for further analysis. However, we retained these low intensities for data analysis if the feature met our filtering criteria. That is, if a feature met our filtering criteria, we still considered intensities belonging to the lowest quartile for data analysis.

We then used these computed intensity quartiles to calculate thresholds for two additional MS1-only datasets. Our first data censoring dataset, “High”, was created similarly to the complete MS1-only dataset, but instead used the lowest intensity of the highest quartile as the missing value threshold (intensity of 29,190,200). Our second data censoring dataset, “High + Medium” instead used the lowest value of the second highest quantile as the missing value threshold (intensity of 12,626,200).

To prevent challenges with the log_2_ transformation and consequent fold change calculations, we then imputed missing data by k-nearest neighbour imputation using the impute R package (Hastie et al., 2019) using default parameters other than colmax=0.95. Feature intensities were then normalized using the median ratio method in the DESeq2 R package (Love et al., 2014). Finally, log_2_ fold change values were calculated for each drug treatment sample using the median value of the control (DMSO) feature. We completed principal component analysis (PCA) on log_2_ fold change values for quality control and to confirm the discriminative abilities of MS1-only identified features.

#### 2.3.2 MS/MS data

Both experiment metadata and MS/MS-level data peptide intensities were imported into R v4.0.4 (R Development Core Team, 2021). Only peptides identified in at least 50% of each drug treatment, including the DMSO control, were kept for further analysis. Peptide intensities were normalized, missing values were imputed and log_2_ fold change values were calculated as previously described. PCA was also completed as described above.

### 2.4 Independent component analysis

The PRECISE ICA implementation was used for robust component calculation (Sastry et al., 2019). Briefly, the PRECISE ICA implementation, originally compiled for RNA-Seq analysis, completes multiple ICA calculations using random seeds. The source components (*S* matrix) identified from each ICA run are clustered, and the final robust components are described as the centroids from each identified cluster. The number of components is set by determining the number of PCA components required to explain 99% variance. Using Python 3.7.5, we called the PRECISE run_ica.py script using default parameters including 100 ICA iterations and a convergence tolerance of 10^−6^.

### 2.5 K-medoid clustering

We used K-medoid clustering to cluster the final *S* matrix using Python v3.7.5 and scikit-learn-extra v0.1.0b2. Clusters were computed from a correlation distance using the “k-medoids+ + + initialization al-gorithm. We tested for cluster fit using *k*=2 to 50 and automated *k* choice by maximum silhouette score. Peptide/feature cluster labels were then exported for further analysis in R.

### 2.6 Eigenfeature-treatment correlation

Using R, eigenfeatures were calculated for each peptide/feature cluster using singular value decomposition (SVD) of each cluster’s intensity values. As described by Langfelder et al. (2007), the first column of SVD is used as the eigenfeature vector for each cluster. These values can then be used to explore relationships between clusters. We calculated Pearson’s correlation coefficients of eigenfeature values to each drug-concentration treatment. Relationships between peptide/feature clusters and drug-concentration treatments were explored by hierarchical clustering. Hierarchical clustering was completed in R using a Pearson’s coefficient correlation distance measure and the average linkage hierarchical clustering method. Robust clusters were identified by two independent methods: 1. Silhouette score calculation and 2. Bootstrapping with 1000 iterations using the pvclust R package (Suzuki et al., 2006) where clusters with approximately unbiased (AU) *p*-values greater than 0.9 were considered robust.

### 2.7 Comparing cluster dendrograms

MS1-only and MS/MS treatment clusters were compared by cophenetic correlation coefficent and visually by tanglegrams. Cophenetic correlation was performed using the `cor_cophenetic()` function from the `dendextend` R package (Galili, 2015). A permutation test using 10,000 iterations was used to estimate the statistical significance of the cophenetic correlation values obtained. In brief, dendrogram labels of the MS/MS acquired clusters were shuffled and cophenetic correlation coefficients were calculated to measure similarity between the MS1-only acquired clusters and the permuted dendrogram. Then, we counted the number of times the absolute value of the permuted correlation coefficient was observed to be greater than the actual calculated value. Tanglegrams of dendrogram similarities were also visualized using `dendextend`.

### 2.8 Data and code availability

Both MS1-only and MS/MS raw data were deposited to the ProteomeXchange Consortium (Deutsch et al., 2017) via the PRIDE (Perez-Riverol et al., 2018) partner repository with the dataset identifiers PXD024815 and PXD024845. Modular and customizable code and example datasets are made available on GitHub at https://github.com/northomics/MetaProClust-MS1. R Notebooks describing the examples in this manuscript and code for figure creation are also included.

## 3 Results

MPC-MS1 is a bioinformatic strategy for rapid screening of MS1 profiles for large scale metaproteomic experiments (Figure 1). In brief, MPC-MS1 uses computationally linked feature intensities acquired from MS1-only mass spectrometry without peptide or protein identification. After feature quantification, a matrix of features by samples, matrix *X*, is decomposed using a robust ICA implementation. The *S* matrix, describing the feature contributions of the original *X* matrix, is then clustered using k-medoids clustering into feature modules. Eigenfeatures, vectors representing a summary of all features of a module, are calculated and correlated with microbiome treatments/conditions. A distance matrix is calculated from these treatment correlations and is used for a final bootstrapped hierarchical clustering of sample treatments. The treatment clusters can then be used to identify groups of treatments or conditions that cause large or small effects on the community of interest in order to prioritize samples for further deep metaproteomic analysis.

**Figure 1:**
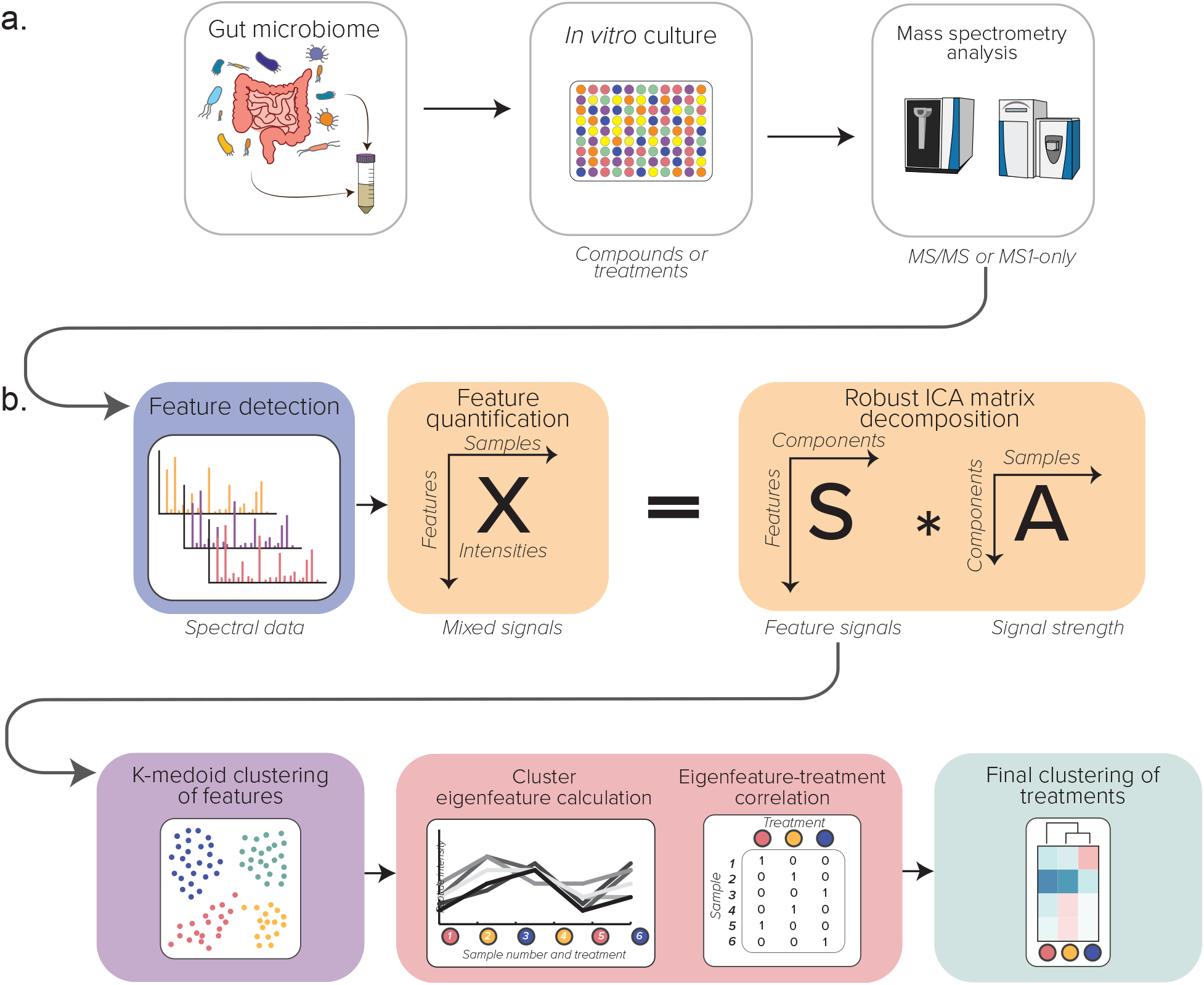
Illustrated schematic of the computational MPC-MS1 workflow. a. The experimental workflow b. The MPC-MS1 framework describing feature detection and quantification from MS1-only spec-tra, ICA matric decomposition, K-medoid feature clustering, eigenfeature calculation and correlation with microbiome treatment and final clustering of microbiome treatments by hierarchical clustering.

As proof-of-concept, we compared MPC-MS1 results of a traditional MS/MS dataset to a dataset composed of rapid MS1-only profiles. Both datasets (MS1-only and MS/MS) were acquired from the same drug treated gut microbiome samples. These metaproteomic samples came from a single gut microbiome treated with five drugs with a variety of known effects on a gut microbiome (Li et al., 2020c) at three concentrations (See Section 2Methods).

### 3.1 MS1-only data

We first completed MPC-MS1 on the MS1-only dataset. First we completed a PCA on log_2_ fold change data to evaluate the discriminative ability of MS1-only features. Overall, drug treatment clustering was observed as is also seen with the MS/MS dataset (Figure 2) PC2 explained drug treatment concentration as a gradient of high to low concentrations loading more positively to negatively on PC2 (Figure S2).

**Figure 2:**
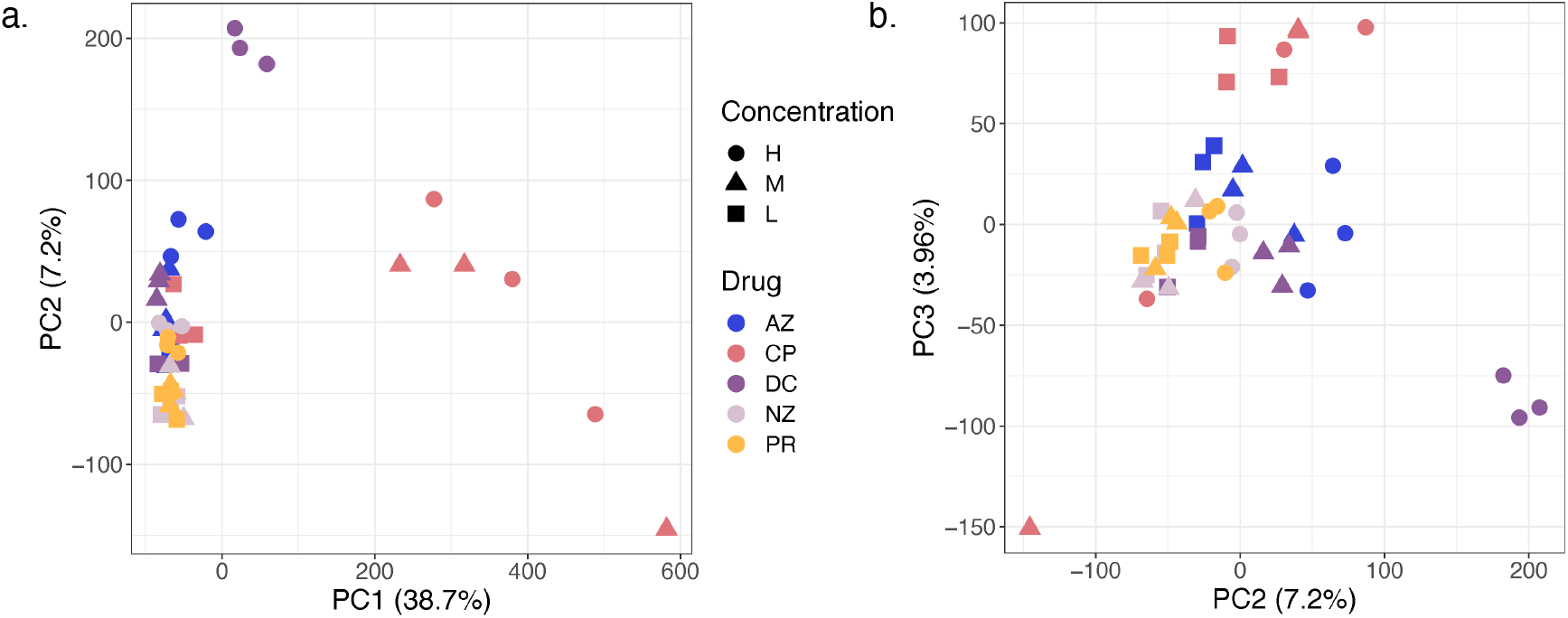
PCA of MS1 log_2_ fold change data. Drug treatments are represented by colours with azathioprine (AZ) in blue, ciprofoxacin (CP) in pink, diclofenac (DC) in light purple, nizatidine (NZ) in dark purple, and paracetamol (PR) in yellow. Drug concentrations are represented by different shapes where high (H) is a circle, medium (M) is a triangle and low (L) is a square. a. Biplot of PC1 vs PC2, b. Biplot of PC2 vs PC3.

PCA was also used to determine the optimal number of components for the consequent ICA process (Sastry et al., 2019) as the number of appropriate ICA components is determined by each data set. In this case, 43 PCA components described at least 99% of the variation in the data. Thus, we completed 100 iterations of ICA matrix decomposition using 43 components. These calculations used 11.36 minutes of CPU time.

The resulting *S* matrix described the contributions of features and was clustered using K-medoid cluster-ing with `k-medoids++` centroid initialization. MPC-MS1 automates *k* or cluster number choice by maximum silhouette score, and in this case *k* = 36 was selected.

Cluster labels were imported into R for eigenfeature calculation using feature intensities. Correlations of cluster eigenfeatures with drug-concentration treatment were calculated using Pearson’s correlation coefficient and used as a distance measure for drug-concentration treatment clustering. Finally, silhouette score metrics identified two large robust drug-concentration clusters (Figure 3). Bootstrapped hierarchical clustering also identified smaller clusters with a AU *p* > 0.9 (Figure S3).

**Figure 3:**
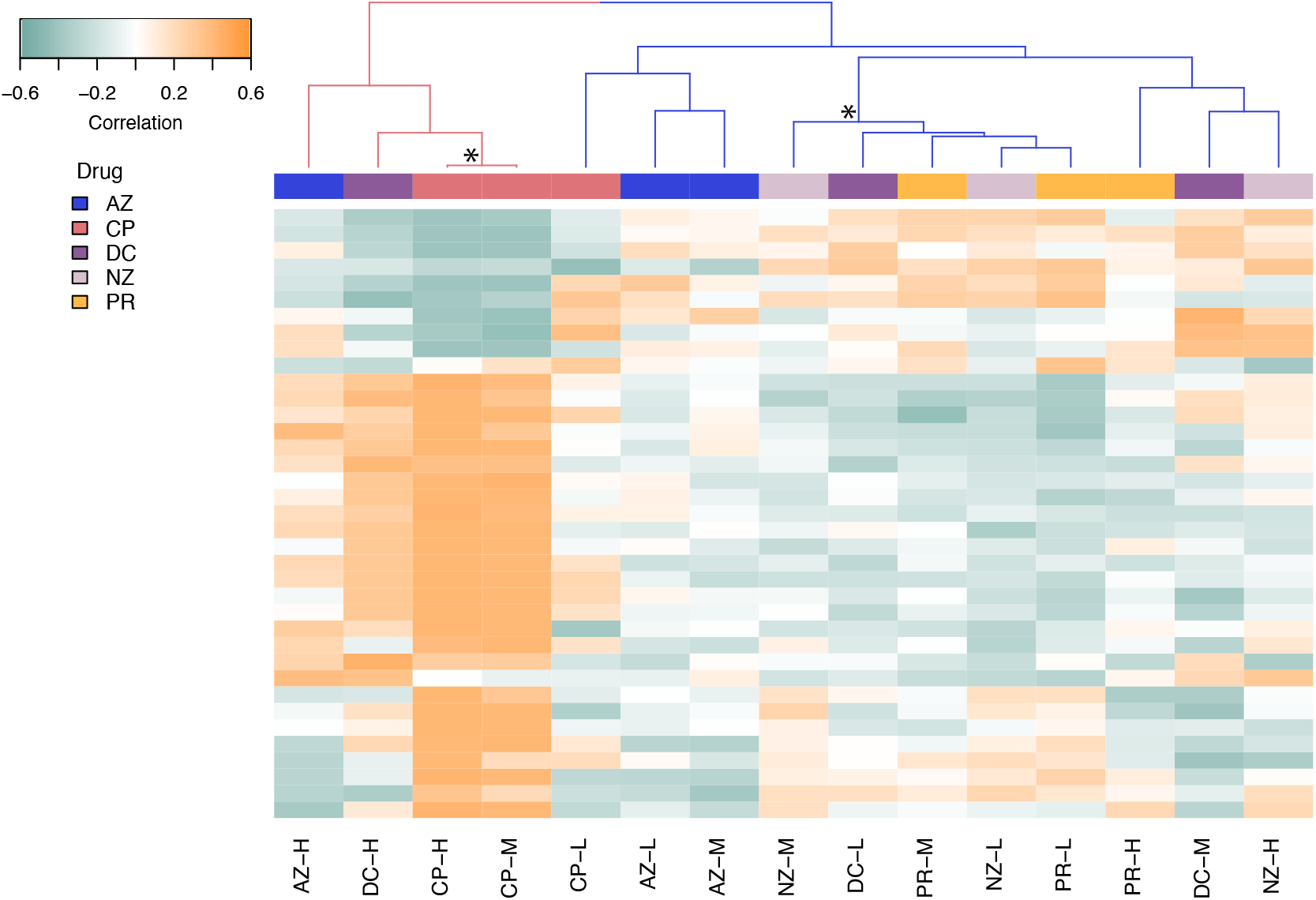
Hierarchical clustering of drug-concentration treatments using the MS1-only dataset. Heatmap rows describe correlation of feature modules with drug-concentration treatments. Blue describes negative correlations while orange describes positive correlations. Using silhouette scores, k=2 was chosen for dendrogram cutting and treatment clusters are coloured as pink and blue. Stars (*) indicate robust nodes as identified in bootstrap analysis (AU *p*-value > 0.9).

The smaller cluster coloured in pink (Figure 3) was dominated by high concentration drug treatments which we expect to have the largest effects on the microbiome. In addition, the pink cluster contains the medium concentration of ciprofloxacin (CP), an antibiotic. The larger cluster coloured in blue contains drug treatments at low (L) and medium (M) concentrations with smaller effects on the microbiome.

#### 3.1.1 MS1-only with data censoring

Using our pre-computed intensity quartiles, we tested if censoring data at intensity level thresholds would change treatment cluster results. In other words, we filtered MS1-only features according to missing values identified by intensity quartiles to create two additional MS1-only datasets: “High” and “High + Medium” intensity feature data. We tested how correlated the clusters inferred from our full MS1 feature dataset would be with the clusters identified by our censored datasets using cophenetic correlation. We showed that both “High” and “High + Medium” datasets produced clusters were correlated with the full MS1 feature dataset (*r* = 0.600 and 0.607 respectively; Figure S4). The “High” and “High + Medium” datasets were, however, significantly highly correlated to each other (*r* = 0.976).

### 3.2 MS/MS data

We again evaluated the discriminative abilities of the classic MS/MS dataset by PCA (Figure 4). In addition, PCA was used to determine the optimal number of components for the consequent ICA process (Sastry et al., 2019) as the number of appropriate ICA components is determined by each data set. PCA was also used to determine the optimal number of components for the consequent ICA process (Sastry et al., 2019) as the number of appropriate ICA components is determined by each data set. In this case, 35 PCA components described at least 99% of the variation in the data. Thus, we completed 100 iterations of ICA matrix decomposition using 35 components which used 13.49 minutes of CPU time.

**Figure 4:**
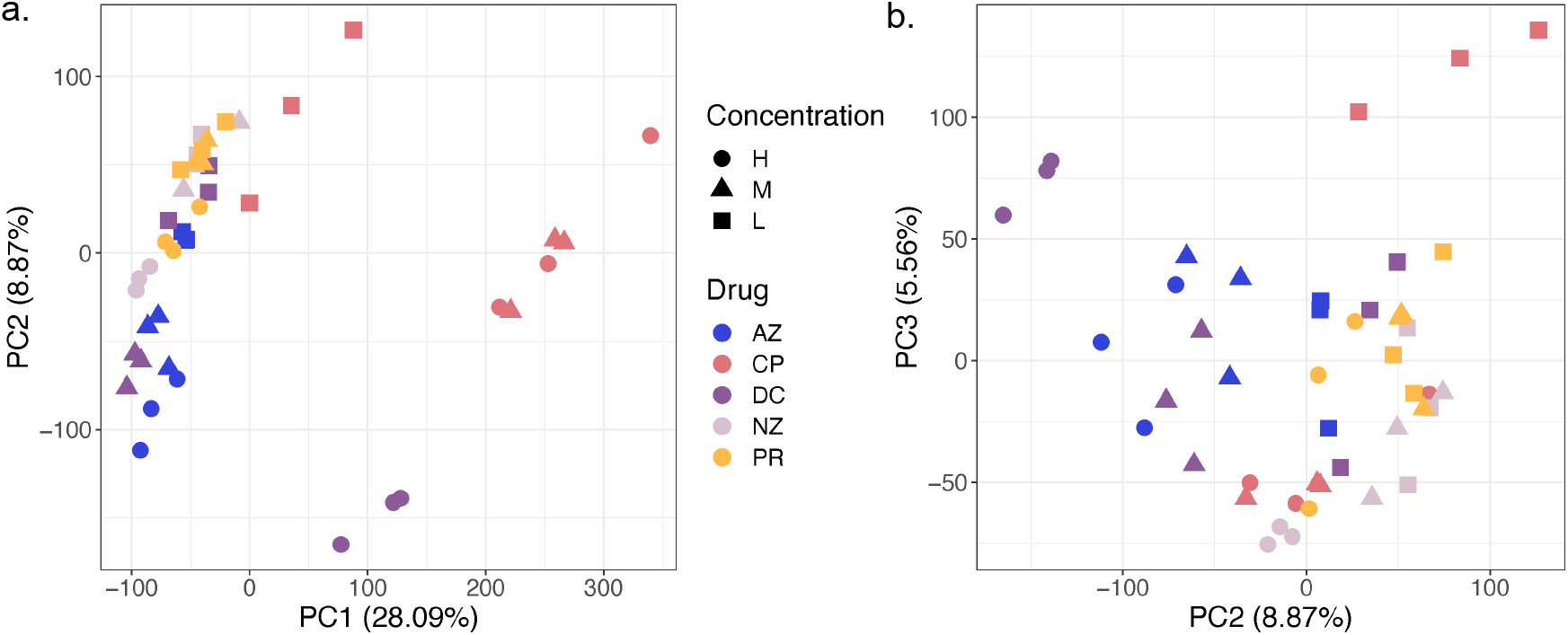
PCA of MS/MS log_2_ fold change data. Drug treatments are represented by colours with azathioprine (AZ) in blue, ciprofoxacin (CP) in pink, diclofenac (DC) in light purple, nizatidine (NZ) in dark purple, and paracetamol (PR) in yellow. Drug concentrations are represented by different shapes where high (H) is a circle, medium (M) is a triangle and low (L) is a square. a. Biplot of PC1 vs PC2, b. Biplot of PC2 vs PC3.

Silhouette scores calculated from k-medoids clustering of the resulting *S* matrix suggested *k* = 48 to be an optimal number of peptide clusters. Peptide cluster labels were then imported into R for eigenfeature calculation and consequent correlation to drug-concentration treatment. Using eigenfeature-treatment cor-relations, we clustered treatments using hierarchical clustering. Silhouette score evaluation identified three robust treatment clusters (Figure 5). Bootstrapped hierarchical clustering also identified the same three clusters as well as four other smaller clusters with bootstrap AU *p*-values > 0.9 (Figure S3). Seven nodes with significant AU *p*-values indicate robust clustering of microbiome treatments.

**Figure 5:**
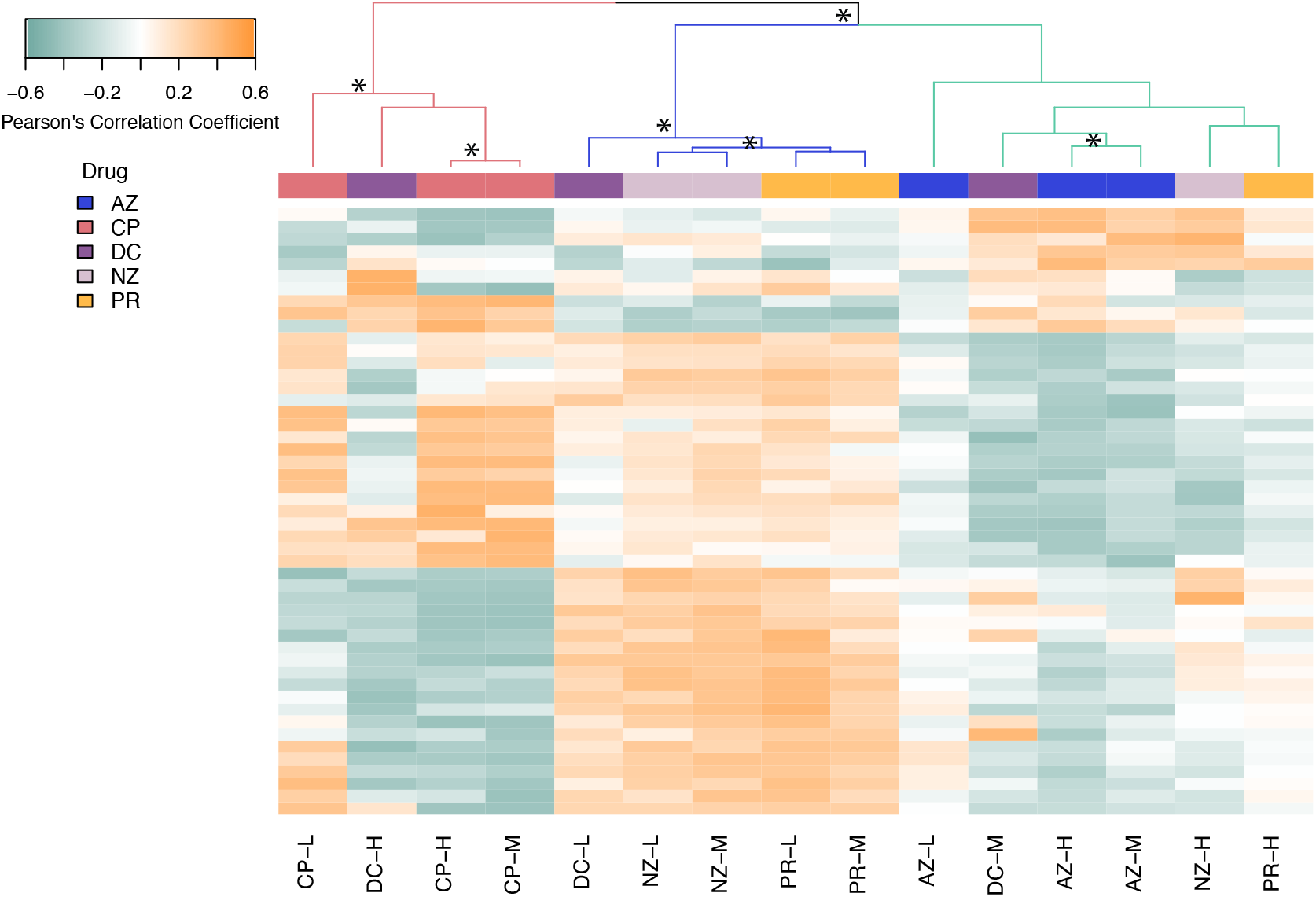
Hierarchical clustering of drug-concentration treatments using the MS/MS dataset. Heatmap rows describe correlation of feature modules with drug-concentration treatments. Blue describes negative correlations while orange describes positive correlations. Using silhouette scores, k=3 was chosen for dendrogram cutting and treatment clusters are coloured as pink, blue and green. Stars (*) indicate robust nodes as identified by bootstrap AU values (AU *p*-value > 0.9).

In general, medium (M) and low (low) concentrations of the same drug tend to cluster together (Figure 5). Medium (M) and low (L) drug concentrations clustering with high (H) concentrations, for example, AZ-L and CP-L, may indicate drugs with larger effects on a gut microbiome even at small concentrations.

### 3.3 Comparing clusters inferred from MS1-only to MS/MS datasets

Finally, we compared clusters inferred from MS1-only feature intensities to peptides identified by canonical MS/MS (Figure 6). The overall topology of both MS1-only and MS/MS inferred clusters remained relatively similar, and only two drug-concentration treatments (CP-L and AZ-H) did not cluster consistently between the two datasets.We confirmed cluster visual similarity through cophenetic correlation calculation of the two dendrograms and found the clusters to be significantly similar and more correlation than the censored data comparisons (*r* = 0.625; permutation *p*-value < 0.0001; Figure S5).

**Figure 6:**
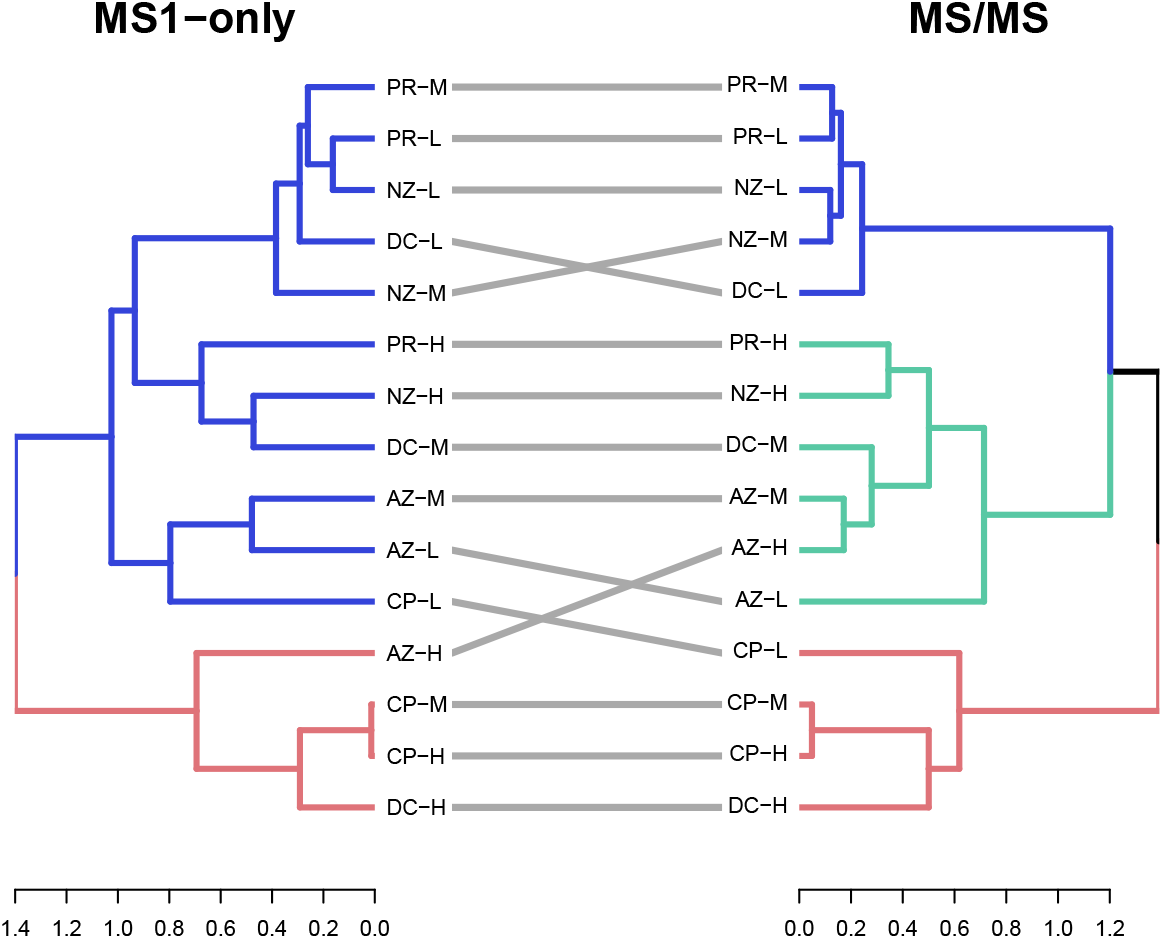
Tanglegram comparing drug-concentration treatment clustering of MS1-only and MS/MS inferred clusters.

## 4 Discussion

While both taxonomically and functionally informative, acquiring MS/MS metaproteomic data for microbiome research can be resource intensive. In particular, data acquisition using the canonical tandem mass spectrometry approach is time consuming, and this challenge is exacerbated in large-scale studies. In this work, we introduce MPC-MS1, a computational tool that uses MS2-independent metaproteomic data and reduces the time and financial requirements typically required in large scale studies. MPC-MS1 identified similar clusters from both MS1-only and MS/MS datasets suggesting that short MS1 gradients are useful for metaproteomic screening. By not relying on peptide identification from MS/MS, MPC-MS1 is not restricted to system-specific databases typically required for confident peptide spectra matches (PSMs) (Tanca et al., 2016).

Rechenberger et al. (2019) recently described the challenges of clinical metaproteomic studies and highlighted that community curated human gut microbiome databases, like IGC, limit peptide identification with over 50% of identified peptides missing from the widely used database. The authors instead suggest sample specific database creation by metagenomics. However, by being MS2-independent, MPC-MS1 removes the challenge of appropriate database selection. MPC-MS1 offers unbiased insights into microbiota community profiles while eliminating the computational and financial requirements of metageonomic database construction and PSM identification. The unrestricted features identified in the MS1-only dataset may also explain the inconsistent treatment clustering observed in the comparison of the MS1-only clusters to those identified using MS/MS (Figure 6) and suggest that features that are only quantified by MS1 play important roles in treatment separation.

Data-dependent acquisition (DDA), a common approach to MS/MS data acquisition in metaproteomics, can also fall short in terms of the high amount of missing values associated with the canonical metaproteomic strategy (O’Brien et al., 2018). For example, DDA only selects peptides with the highest intensities measured by MS1 for further fragmentation and eventual identification by MS2. Missing values can occur when a peptide is not selected for MS2 analysis, thus biasing MS/MS experiments to higher intensity peptides. As described by Ivanov et al. (2020), there is developing interest in MS2-independent proteomics to remove the challenge of these technically imposed missing values and increase reproducibility in large scale proteomics studies.

While MS2-independent proteomic studies are of interest and seem promising there are very few studies available to have used the approach. DirectMS1 is an example of a short MS1 gradient tool for protein identification and quantification that was developed and tested on HeLa cells (Ivanov et al., 2020). Alternatively, RIPPER is an MS1-based proteomic and metabolomic tool that uses MS1 features without protein identi-fication (Van Riper et al., 2016). RIPPER’s main goal is to use t-tests to identify quantitative differences between two treatments. To our knowledge, MPC-MS1 is the first bioinformatic tool that considers MS1 profiling of metaproteomic data, and the first MS1-only tool to accept more than two treatment conditions in its analysis. Because MPC-MS1 was developed for metaproteomics, MPC-MS1 is currently limited to treatment separation and does not identify peptides. However,in the future MPC-MS1 can be extended for protein identification through MS1 features using peptide mass fingerprinting (James et al., 1993; Pappin et al., 1993; Mann et al., 1993; Yeats et al., 1993) and an MS1-only search workflow (Ivanov et al., 2017). However, this approach remains to be tested on complex metaproteomic data and is currently beyond the scope of our study.

Methods other than MS2-independent proteomics have also been suggested for the reduction of required resources for proteomic studies. For example, Meyer et al. (2020) maintain MS/MS and instead remove liquid chromatography (LC) to perform data independent acquisition (DIA) proteomics by direct infusion with the use of high-field asymmetric waveform ion mobility spectrometry (FAIMS). While FAIMS, and ion mobility in general, presents an intriguing venue in proteomics, the complexity of microbiome metaproteome samples may challenge its utility. For instance, the compensation voltage must be either kept fixed or scanned across a narrow range to prevent the excessive prolongation of the duty cycle (Creese et al., 2013). Therefore, DIA via direct infusion and FAIMS may not be an appropriate approach for metaproteomic samples. However, in our proof of concept study, we demonstrated the success of not only rapid metaproteomic screening for microbiome treatments, but also the possibility to use short MS1 gradients in microbiome studies. To note, the reasonably high cophenetic correlation of “High” intensity MS1 feature clusters with the full MS1-only feature dataset also suggests there is a possibility for further reduction in MS1 gradient time (Figure S4). While we had originally tested data censoring of MS1 features due to the inherent inclusion of noise in MS1-only data, the reduction in cluster similarity to those inferred from the full feature dataset suggests that the so-called “noise” of MS1-only data could be essential to treatment separation.

The framework for MPC-MS1 was inspired by gene expression network analysis tools commonly used in transcriptomics. A recent meta-analysis explored multiple module detection techniques in this type of network analysis and evaluated the performance of each algorithm by comparing detected to known modules (Saelens et al., 2018). Matrix decomposition methods, such as ICA, consistently achieved high scores in multiple evaluation methods, and previous research has used ICA to cluster bladder cancer subtypes (Biton et al., 2014) and to identify robust gene modules in *Escherichia coli* (Sastry et al., 2019). ICA is often described as a “blind source separation” linear transformation method that is used to identify a linear representation of independent sources in set of mixed signals (Cherry, 1953). A common example using blind source separation is described as the “cocktail party problem” where one attempts to determine what a person is saying in a noisy room, such as at a cocktail party (Cherry, 1953) Similar to a “cocktail party”, one can use ICA to separate non-biological or individualistic factors observed in microbiome data using this matrix decomposition method. MPC-MS1 uses ICA to identify robust modules of quantified features that are then used to cluster microbiome treatments.

The main intended use of MPC-MS1 is as a screening tool for more efficient, resource conscious and intentional metaproteomic research. In its current implementation, MPC-MS1 identifies feature modules that summarize MS1 profiles for treatment clustering. However, these modules also contain features with similar intensity profiles throughout an entire experiment that may warrant further investigation themselves. For example, gut microbiome clinical research may be interested in MS1-only biomarker discovery. If a module is significantly correlated with a microbiome condition of interest, that module may contain candidate biomarkers. In this study, we used MPC-MS1 to screen for compounds with effects on the gut microbiome, and we hope MPC-MS1 will also be used in clinical studies. In addition, it is possible to use MPC-MS1 for more efficient MS/MS analysis. For example, novel treatments or conditions may cluster with known microbiome perturbing treatments and can be selected for reanalysis by deep metaproteomics.

The MPC-MS1 computational pipeline is an effective bioinformatic tool for metaproteomic screening that can be used on any operating system that supports R and Python. By using rapid MS1 profiles rather than time consuming tandem mass spectrometry, MPC-MS1 reduces the resources typically required for deep metaproteomic experiments. Our tool acts in compliment to large scale RapidAIM assays and avoids the challenge of database selection and peptide identification because it is an MS2-independent approach. By removing peptide identification, MPC-MS1 does not require a protein database or a false discovery rate reducing search strategy. Instead, MPC-MS1 is a rapid and unbiased tool that can be used to screen large scale metaproteomic experiments and may be especially relevant to clinical studies. Finally, to our knowledge, MPC-MS1 is the first MS1 profiling tool specifically developed for the metaproteomic community.

## 5 Acknowledgements

This work was funded by the Government of Canada through Genome Canada and the Ontario Genomics Institute (OGI-156 and OGI-149), the Natural Sciences and Engineering Research Council of Canada (NSERC, grant no. 210034), and the Ontario Ministry of Economic Development and Innovation (ORF-DIG-14405 and project 13440) as well as NSERC Discovery Grant to M.L.A. C.M.A.S. was funded by a stipend from the NSERC CREATE in Technologies for Microbiome Science and Engineering (TECHNOMISE) Program.

## 6 Author Contributions

Conceptualization, C.M.A.S. and Z.N.; Methodology, C.M.A.S. and Z.N.; Software, C.M.A.S; Investigation, C.M.A.S., Z.N., L.L. and M.M.K.; Formal Analysis, C.M.A.S.; Writing - Original Draft, C.M.A.S. Writing- Review & Editing, C.M.A.S., Z.N., L.L., M.M.K., X.X., M.L.A. and D.F.; Resources, D.F.; Supervision, M.L.A. and D.F.; Funding Acquisition, D.F.

---

**Figure S1:**
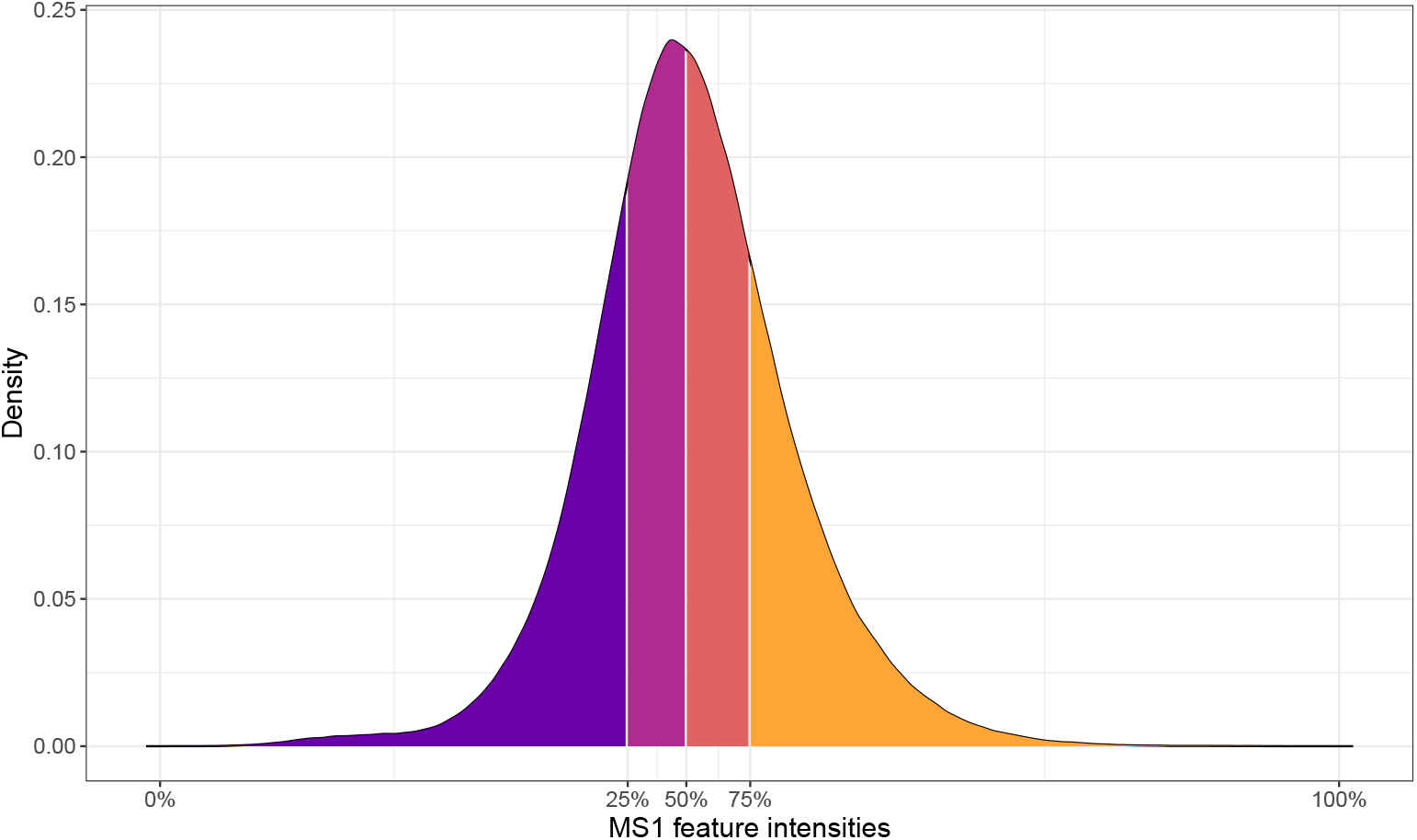
Density of MS1-only feature intensities. Quartiles are indicated by colours and are separated by white lines. The percent is also indicated on the x-axis.

**Figure S2:**
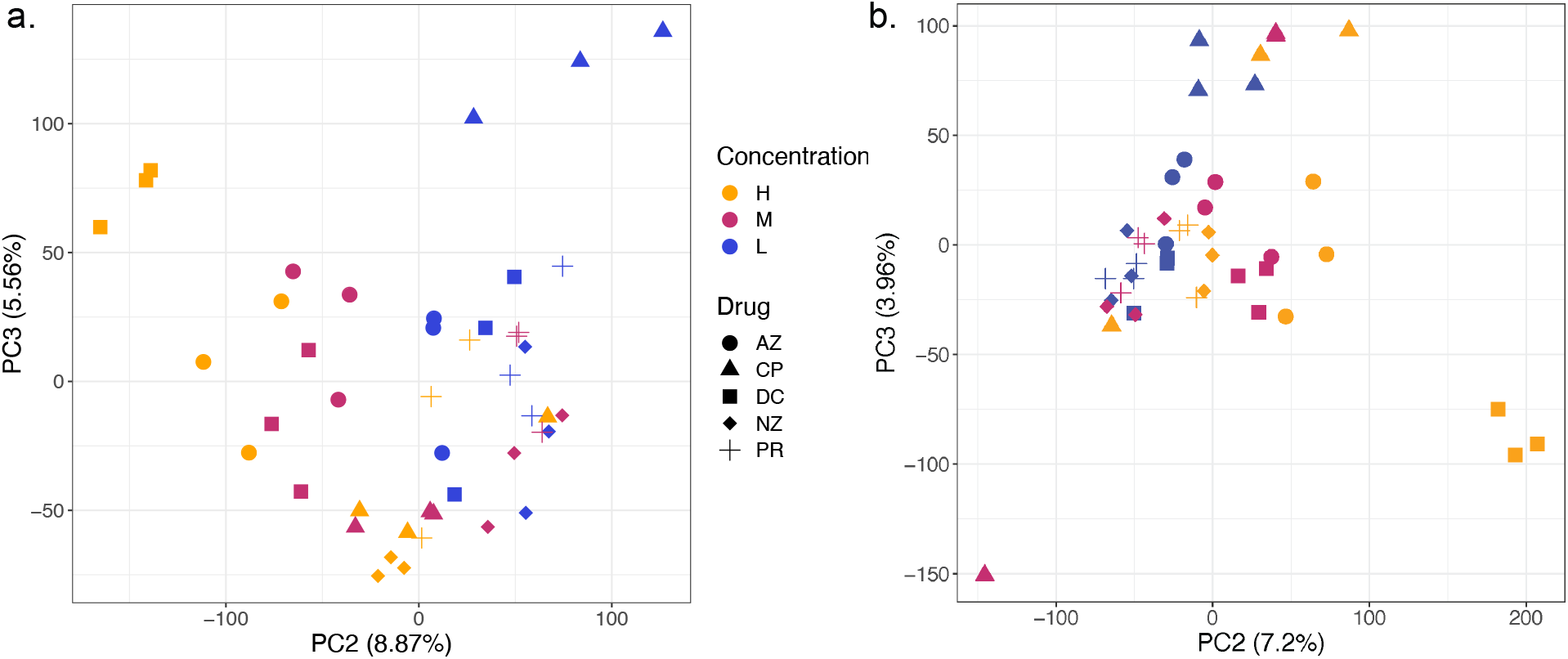
PCA of MS/MS and MS1 only fold change data. Plot is coloured using drug concentrations to highlight the gradient observed on PC2. a. Classic MS/MS acquired data, b. MS1-only data.

**Figure S3:**
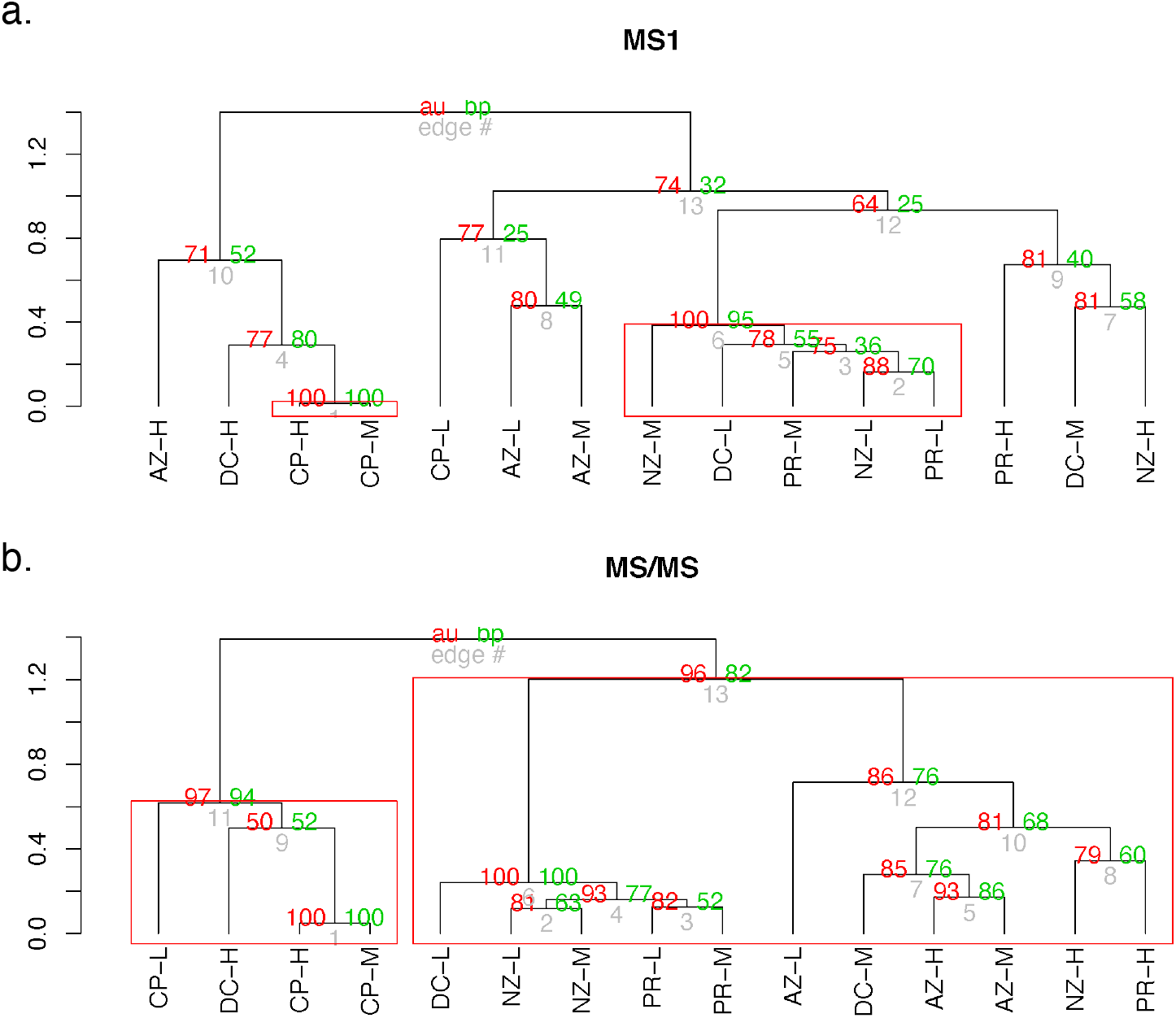
Bootstrapped hierarchical clustering dendrograms for MS1-only and MS/MS datasets. Approximately unbiased (AU) P-values were calculated using 1000 iteractions of multiscale bootstrap resampling. Red values at nodes describe approximately unbiased (AU) p-values. Green values at nodes describe Bootstrap Probability (BP) values but were not used in the evaluation of clusters. a. describes MS1-only hierarchical clustering, b. describes MS/MS hierarchical clustering.

**Figure S4:**
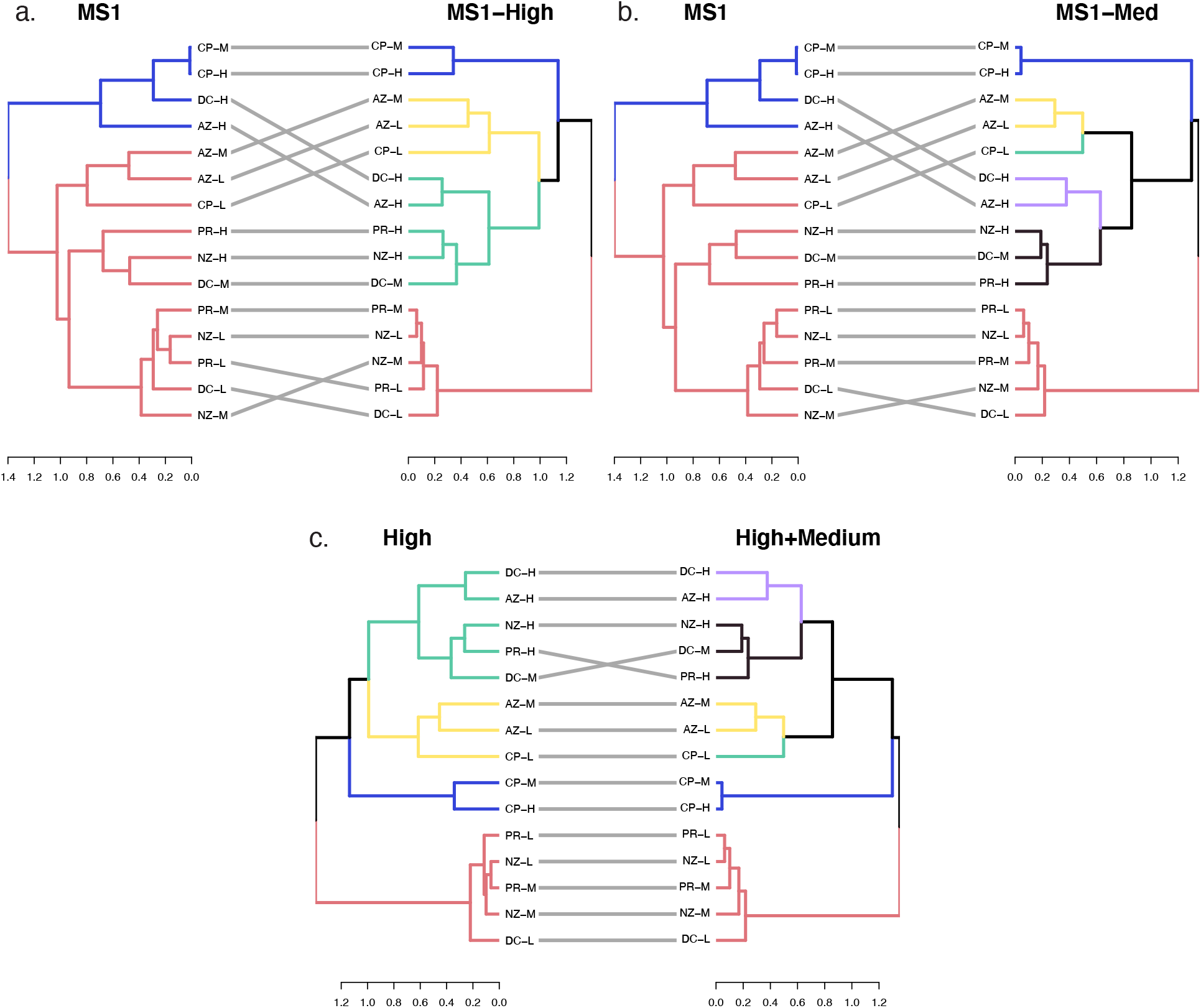
Tanglegrams of clusters identified by data censoring. Tanglegrams were rotated to reduce entanglement measures using the ‘step2side’ untangle function in the R package. a. Final MS1-only clusters compared to high intensity MS1 features, b. final MS1-only clusters compared to high and medium intensity features, c. High intensity clusters compared to high and low intensity features.

**Figure S5:**
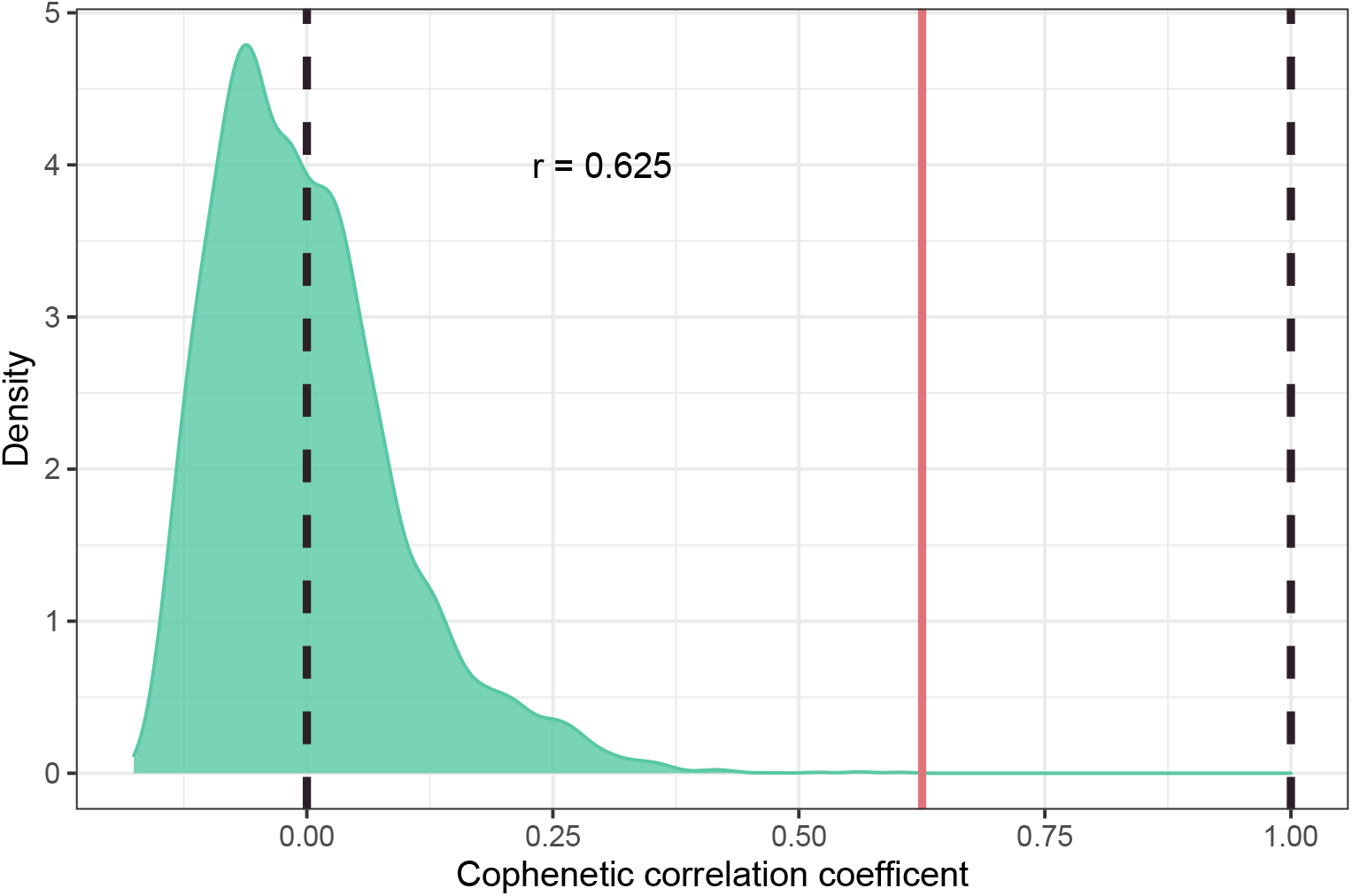
Distribution of permuted cophenetic correlation coefficient comparing MS1-only to MS/MS dendrograms. Permuted cophenetic correlation coefficients were computeted for 2000 iterations by dendrogram label swapping. The permutation *p*-value < 0.0001 was calculated by the total number of absolute values of perumuted corelation coefficients that was greater than the calculated cophenetic coefficient (*r* = 0.625).

## Notes

https://github.com/northomics/MetaProClust-MS1

### Competing Interest Statement

D.F. co-founded MedBiome, a clinical microbiomics company. The remaining authors declare no competing interests.

